# Evaluating valid parameter regimes for biocircuits

**DOI:** 10.64898/2026.01.19.700491

**Authors:** Qinguo Liu, Xinying Ren, Fangzhou Xiao

## Abstract

Biocircuit functions are often valid only in specific parameter regimes, yet these regimes are rarely made explicit. We use a holistic analysis method based on regimes to derive validity conditions and introduce the Realizability Index (*R*-index), quantifying the size of the valid regions in log-parameter space. The framework is applied to Michaelis-Menten kinetics, Hill functions, and enzymatic negative-feedback adaptation, showing how circuit structure and experimental control variables shape functional realizability. Our analysis shows the Hill function’s *R*-index goes to zero in sequential binding with increasing Hill coefficient. We also resolve an active debate about whether negative-feedback adaptation is realizable when competitive binding is taken into account, and demonstrates the superiority of the holistic *R*-index method over numerical parameter scans that lead to incorrect conclusions. *R*-index defines a validity-aware language for studying and designing functional biocircuits.

## 1. INTRODUCTION

Mechanistic chemical reaction networks are a central language for modeling biomolecular information processing in systems and synthetic biology (Alon, 2019). They are mechanistically faithful, but their many states and poorly known parameters make general analysis and rational design difficult. For this reason, simplified descriptions such as Michaelis-Menten kinetics, Hill functions, and reduced feedback models are often written down directly without full justification.

These approximations are useful only in appropriate regimes. If their validity conditions are not checked, a designed biocircuit may satisfy the reduced model but fail experimentally. This problem is especially acute when protein abundances, binding constants, and catalytic rates vary over orders of magnitude and are only partially controllable.

Reaction Order Polyhedra (ROP) theory provides a way to analyze biochemical networks without first assuming a particular approximation (Marken et al., 2020; Xiao et al., 2021; Xiao, 2022). ROP partitions log-concentration and log-parameter space into dominance regimes, within which the network admits simple asymptotic descriptions. Here we use this regime-based view to derive validity conditions for desired functions and to quantify their size by a Realizability Index (*R*-index).

The paper proceeds as follows. Section 2 defines the *R*-index and summarizes the regime-based computation. Section 3 applies the method to Michaelis-Menten and Hill approximations. Section 4 applies it to adaptation in a competitive enzymatic negative-feedback biocircuit.

## 2. DEFINITION OF THE *R*-INDEX AND REGIME ANALYSIS

Consider a desired function of a biocircuit with free positive parameters *x*_1_, …, *x*_*n*_, including concentrations and constants of binding or catalysis. In log space, write *z*_*i*_ = log *x*_*i*_. A **regime polyhedron** is described by linear inequalities

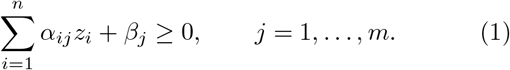

The corresponding asymptotic **regime cone** is obtained by dropping the constant offsets,

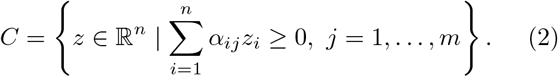

We define the *R***-index** of the regime as the solid-angle fraction

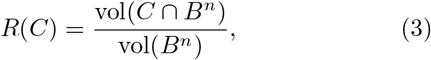

where *B*^*n*^ is the unit ball. Numerically, we estimate this fraction by Gaussian sampling. For a union of non-overlapping valid regimes, the *R*-index is the sum of their fractions.

Table 1 and Fig. 1 give simple examples, where asymptotic inequalities are treated as inequalities when calculating *R*-index. A single inequality *A* ≫ *B* occupies half of log-parameter space. Two inequalities requiring *A* to be the largest of *A, B, C* occupy one third. A strict ordering *A* ≫ *B* ≫ *C* occupies one of the 3! possible orderings. By contrast, an approximate equality *A* ≈ *B* defines a finite-width band around a hyperplane, so its asymptotic volume fraction is zero unless the equality is enforced structurally and the two variables are not independent.

**Table 1.** *R*-index examples.

| Condition | Interpretation | $R$-index |
| --- | --- | --- |
| $A \gg B$ | one ordering of two variables | 1/2 |
| $A \gg B, A \gg C$ | $A$ is largest among three | 1/3 |
| $A \gg B \gg C$ | one complete ordering over three | 1/6 |
| $A \approx B$ | finite-width envelope | 0 |

**Fig. 1.**
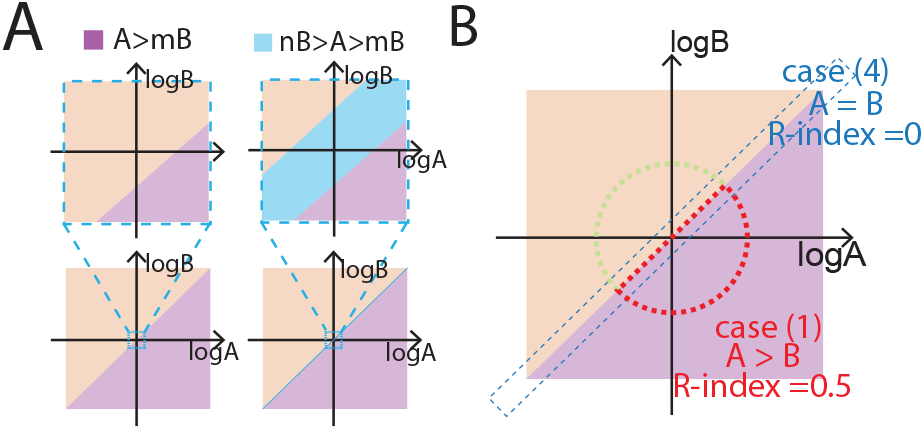
Geometric interpretation of the *R*-index for an inequality regime and an equality-like regime.

For binding networks, the regimes are obtained from the binding equilibria and conservation laws. For the elementary reactions in one enzyme-substrate binding:

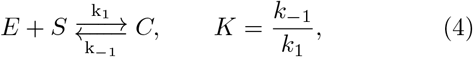

the binding equilibria and conservation laws give

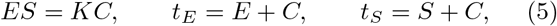

where *t*_*E*_ and *t*_*S*_ are total enzymes and substrates, the conserved quantities of the binding network.

We would like to analyze the regimes of this network in terms of (*t*_*E*_, *t*_*S*_, *K*) as the natural **parameter space**, since production and degradation of molecules change the totals, while different molecules vary the binding constants.

The dominance choices in the two conservations define four regimes. Namely, *t*_*E*_ could be dominated by either *E* or *C*, and *t*_*S*_ could be dominated by either *S* or *C*. Solving the equilibrium equations inside each dominance pattern maps the species-space regimes to conditions in the natural parameter space (*t*_*E*_, *t*_*S*_, *K*).

For example, Eq. (5) becomes the following conditions under regime (*t*_*E*_, *t*_*S*_) ≈ (*E, S*):

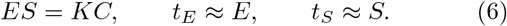

In particular, the dominance conditions specify the following asymptotic inequalities in (*E, S, C*), or **species space**:

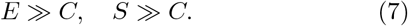

This maps to asymptotic inequalities in totals and binding constants (*t*_*E*_, *t*_*S*_, *K*), or **parameter space**, via Eq. (5):

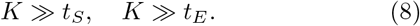

Treating asymptotic conditions as inequalities yield a regime polyhedron:

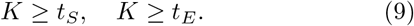

This produces the parameter-space *R*-indices listed in Table 2; the fourth case has zero *R*-index because it requires an equality.

**Table 2.** Regimes of the elementary binding reaction in (*t*_*E*_, *t*_*S*_, *K*) space.

| Regime | Dominance | Parameter condition | $R$-index |
| --- | --- | --- | --- |
| #1 | $(E, S)$ | $K \gg t_S, K \gg t_E$ | 1/3 |
| #2 | $(E, C)$ | $t_E \gg K, t_E \gg t_S$ | 1/3 |
| #3 | $(C, S)$ | $t_S \gg K, t_S \gg t_E$ | 1/3 |
| #4 | $(C, C)$ | $t_E \approx t_S, t_E \gg K$ | 0 |

Fig. 2 illustrates how each dominance regime in species space map to parameter space. Note that *R*-index in species space could be drastically different from that in parameter space. For example, *R*-index of Regime #4 is 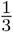 in species space, but 0 in parameter space, while that of Regime #1 is 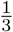 in both spaces.

**Fig. 2.**
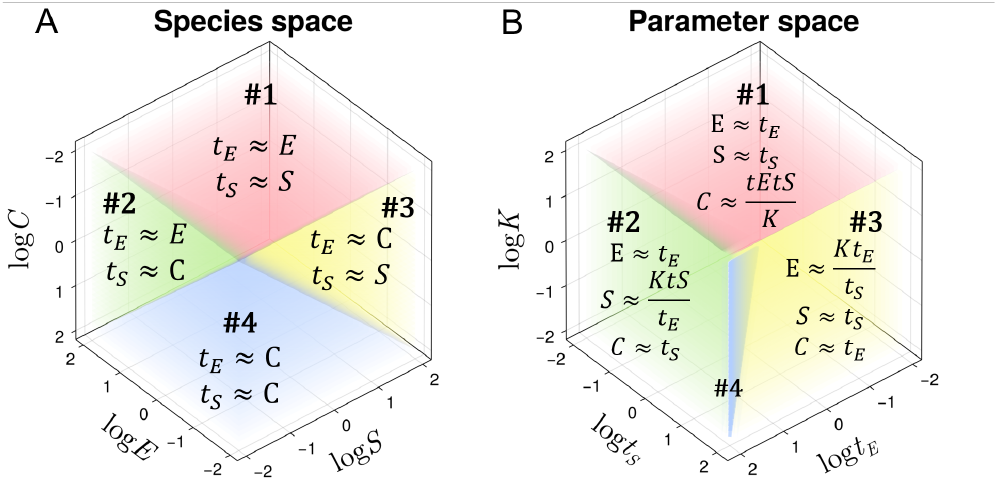
Regime partition for the elementary binding reaction in (A) species(*E, S, C*) space and (B) parameter(*t*_*E*_, *t*_*S*_, *K*) space.

The same procedure extends to larger binding networks. In practice, our computation has four steps. First, enumerate all dominance choices in each conservation equation. Second, solve the binding equilibrium equations under those dominance choices to express species concentrations in terms of total concentrations and constants. Third, substitute the solution back into the dominance inequalities to obtain the regime polyhedron in the chosen parameter space. Finally, find which regimes have the desired functional properties and estimate the solid-angle fraction of their union as the function’s *R*-index.

Note that the same function in a biochemical network can have different *R*-indices depending on which variables are externally controlled and which variables must vary freely during operation.

A regime with positive *R*-index should be interpreted as structurally realizable rather than automatically easy to implement. Experimental bounds, correlations between parameters, or parameter ranges inherited from a previous design can increase or decrease the fraction. The unconstrained *R*-index therefore serves as a baseline structural score, holistic and unbiased by priors from specific scenarios. On top of this, constrained versions can then be computed by sampling from the experimentally implementable or biologically relevant region.

The source code used to compute regimes and their *R*-indices is available at https://github.com/Qinguo25/BindingAndCatalysis.jl/tree/ifac2026.

## 3. *R*-INDEX OF MICHAELIS-MENTEN AND HILL FUNCTIONS

### 3.1 Michaelis-Menten kinetics

Consider the enzymatic reaction

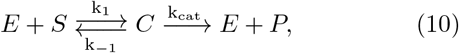

with total concentrations *t*_*E*_ = *E* + *C* and *t*_*S*_ = *S* + *C*.

To analyze how the catalysis rate *k*_cat_*C* changes with total concentrations, two separate approximations are often made. First, the total quasi-steady-state approximation assumes that the concentration of *C* relaxes rapidly relative to catalysis progress, so *ES* = *K*_*m*_*C* holds on the slow catalytic timescale, where *K*_*m*_ = (*k*_*cat*_+*k*_−1_)*/k*_1_. This approximation is shown to roughly always hold (Tzafriri, 2003). Second, most substrates are in free form, so *t*_*S*_ ≈ *S*. This yields the renowned Michaelis-Menten expression

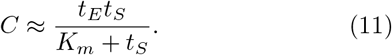

A common sufficient condition for this is *t*_*S*_ ≫ *t*_*E*_, because *C* ≤ *t*_*E*_ then implies *t*_*S*_ = *S* + *C* ≈ *S*. This condition is simple but not necessary, and it can be violated in cellular contexts where both enzyme and substrate are macromolecules with comparable abundance.

In Fig. 3, we compare the exact complex concentration with the Michaelis-Menten formula, showing large errors in certain regions. We now use *R*-index to holistically evaluate the valid regimes of the Michaelis-Menten formula.

**Fig. 3.**
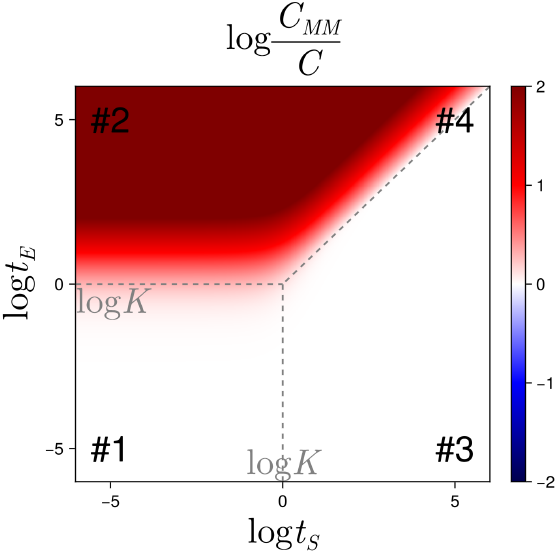
Comparison of the exact solution for *C* with the Michaelis-Menten approximation *C*_*MM*_.

The steady state and conservation equations of this system are the same as Eq. (5), with *K* replaced by *K*_*m*_. So regime analysis of Eq. (5) directly applies here as well. Regimes #1 and #3 in Table 2 have *t*_*S*_ ≈ *S* and together reproduce the two Michaelis-Menten limits *C* ≈ *t*_*E*_*t*_*S*_*/K*_*m*_ and *C* ≈ *t*_*E*_. Therefore, treating (*t*_*E*_, *t*_*S*_, *K*_*m*_) as controllable parameters, the validity condition becomes

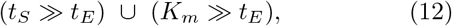

consistent with the result reported in (Schnell and Mendoza, 1997). The corresponding *R*-index is 2*/*3. This is larger than the sufficient condition *t*_*S*_ ≫ *t*_*E*_ usually used to derive Michaelis-Menten, whose *R*-index is only 1*/*2.

The *R*-index changes with the control variables. If the model is used over a full dynamic range of *t*_*S*_ while only (*t*_*E*_, *K*_*m*_) are design parameters, the validity condition reduces to *K*_*m*_ ≫ *t*_*E*_, giving *R* = 1*/*2. Thus the *R*-index should always be interpreted relative to the variables that are independently tunable in the experiment. Fig. 4 shows how the validity region changes when *t*_*S*_ is free to vary.

**Fig. 4.**
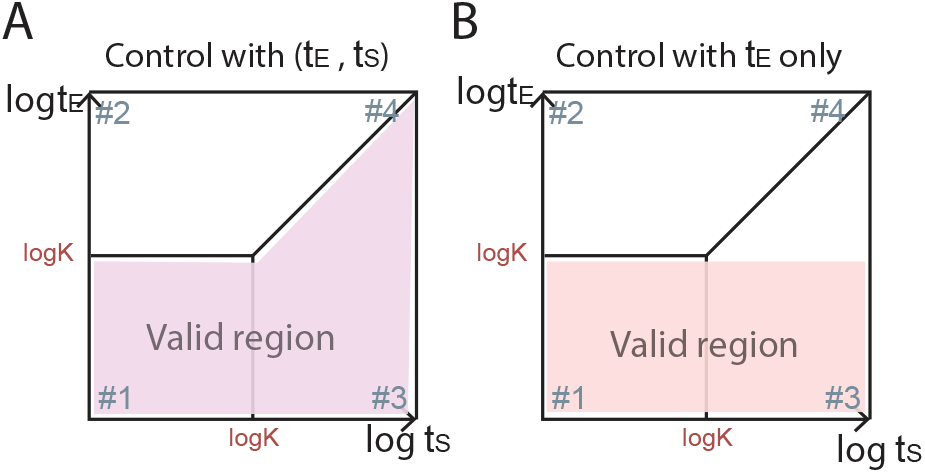
Validity regions for Michaelis-Menten kinetics under different choices of control variables.

### 3.2 Hill functions

The Hill form

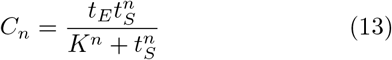

is often used for cooperative binding, and *n* is the number of substrate binding sites the enzyme has.

In terms of regimes, the Hill function prescribes a regime sequence where 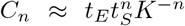 holds for all regimes feasible with *t*_*S*_ ≪ *K*, i.e. containing points satisfying *t*_*S*_ ≪ *K*, and *C*_*n*_ ≈ *t*_*E*_ holds for all regimes feasible with *t*_*S*_ ≫ *K*.

Let us start with *n* = 2. We analyze two mechanisms: sequential binding and dimer binding, shown in Fig. 5 and Table. 3.

**Fig. 5.**
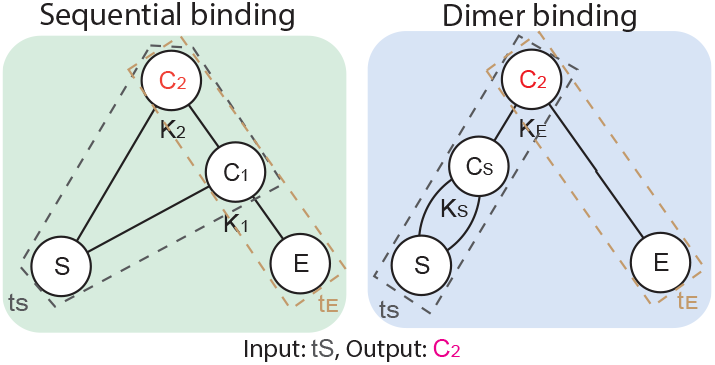
Sequential and dimer mechanisms that can both lead to an *n* = 2 Hill-like expression in selected regimes.

**Table 3.**
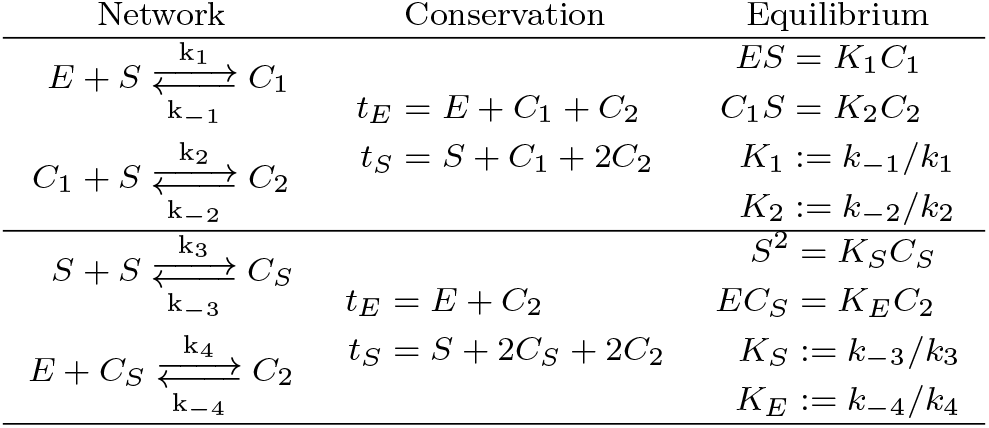
The binding networks and equations for the sequential binding network (top) and the dimer binding network (bottom)

The sequential binding network with *n* = 2 has eight regimes. The only sequence of regimes that fits the Hill prescription is #1:(*t*_*E*_, *t*_*S*_) ≈ (*E, S*) and #7:(*C*_2_, *S*). This is numerically validated in Fig. 6A. Hill form’s *R*-index is 0.56 when we treat (*t*_*S*_, *t*_*E*_, *K*_1_, *K*_2_) as parameters, and it shrinks to 0.25 when the parameters are (*t*_*E*_, *K*_1_, *K*_2_), so that *t*_*S*_ can take the full dynamic range.

**Fig. 6.**
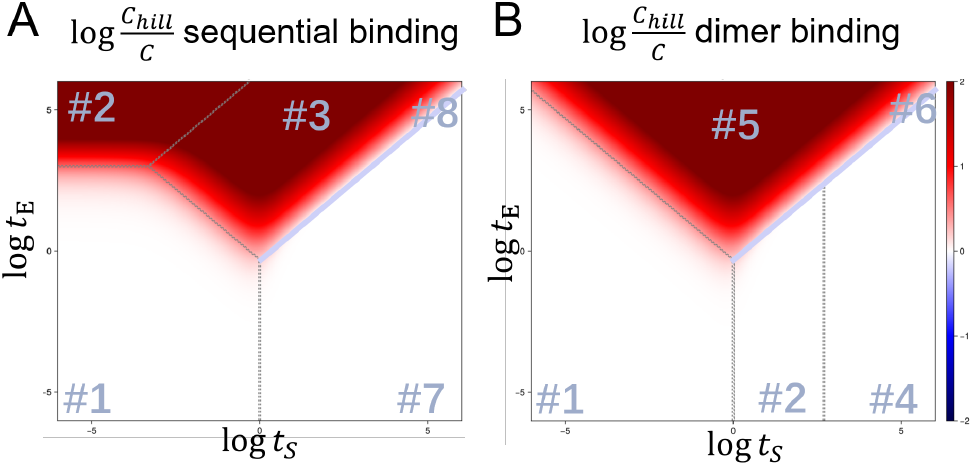
Numeric comparison between exact solved *C* with Hill function derived *C*, denoted as *C*_*hill*_. (A) Sequential binding network with *n* = 2, slice with log_10_ *K*_1_ = 3, log_10_ *K*_2_ = −3. The regimes are indexed by #1:(*t*_*E*_, *t*_*S*_) ≈ (*E, S*), #2:(*E, C*_1_), #3:(*E, C*_2_), #4:(*C*_1_, *S*), #5:(*C*_1_, *C*_1_), #6:(*C*_1_, *C*_2_), #7:(*C*_2_, *S*), and #8:(*C*_2_, *C*_2_). (B) Dimer binding network slice with log_10_ *K*_*E*_ = −3, log_10_ *K*_*S*_ = 3. The regimes are indexed by #1:(*t*_*E*_, *t*_*S*_) ≈ (*E, S*), #2:(*C*_2_, *S*), #3:(*E, C*_*S*_), #4:(*C*_2_, *C*_*S*_), #5:(*E, C*_2_), and #6:(*C*_2_, *C*_2_).

The dimer binding network with *n* = 2 has six regimes. The only sequence of regimes that fits the Hill prescription is #1:(*E, S*), #2:(*C*_2_, *S*), and #4:(*C*_2_, *C*_*S*_), where #2 and #4 share the same expression *C*_2_ = *t*_*E*_. This is numerically validated in Fig. 6B. Hill form’s *R*-index is 0.646 when we treat (*t*_*S*_, *t*_*E*_, *K*_*E*_, *K*_*S*_) as parameters, and it shrinks to 0.25 when the parameters are (*t*_*E*_, *K*_1_, *K*_2_), so that *t*_*S*_ can take the full dynamic range.

For sequential binding with generic *n*, we find the Hill regime sequence generalizes the *n* = 2 case: regimes (*E, S*) and (*C*_*n*_, *S*). Taking (*t*_*S*_, *t*_*E*_, *K*_1_, …, *K*_*n*_) as parameters, the numerically observed sequence (in Fig. 7)is well fit by

**Fig. 7.**
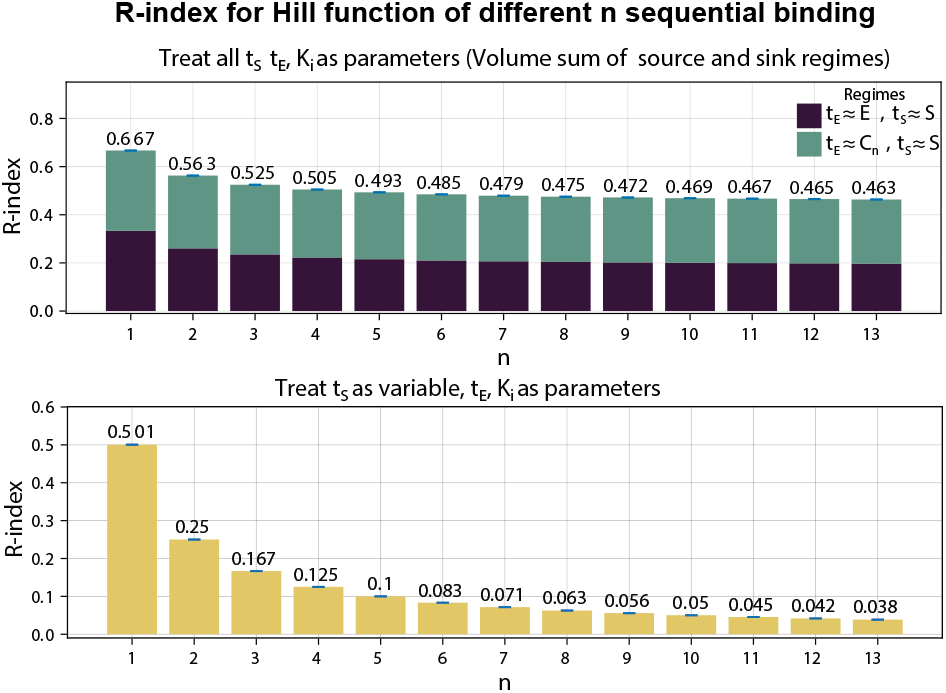
The *R*-index of sequential Hill functions with *n* substrates when *t*_*S*_ is treated as a parameter (top) or as a free variable (bottom).

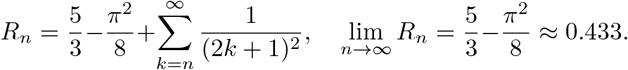

If *t*_*S*_ takes the full dynamic range therefore removed from the parameters, then the *R*-index decays approximately as 1*/*(2*n*). Importantly, it approaches zero for large *n*. This explains why a high-*n* Hill fit can be locally useful while being structurally fragile as a global reduced model.

The results of this section are summarized in Table 4.

**Table 4.** Summary of Hill-function *R*-indices. “All” treats *t*_*S*_ as a parameter; “free *t*_*S*_” requires validity over the *t*_*S*_ range.

| Model | $R_{\text{all}}$ | $R_{\text{free}}$ |
| --- | --- | --- |
| MM ($n = 1$) | 0.667 | 0.500 |
| Seq., $n = 2$ | 0.562 | 0.250 |
| Dimer, $n = 2$ | 0.646 | 0.250 |
| Seq., $n$ | $\frac{5}{3} - \frac{\pi^2}{8} + \sum_{k=n}^{\infty} \frac{1}{(2k+1)^2}$ | $\frac{1}{2n}$ |

## 4. *R*-INDEX OF ADAPTATION IN A NEGATIVE FEEDBACK BIOCIRCUIT

Adaptation means that a step input produces a transient response but the output returns to its pre-stimulus steady-state level. We study the enzymatic negative-feedback loop of (Ma et al., 2009), shown in Fig. 8. The input is *t*_*I*_ and the output is either the free active species *A*^∗^ or the total active species 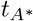.

**Fig. 8.**
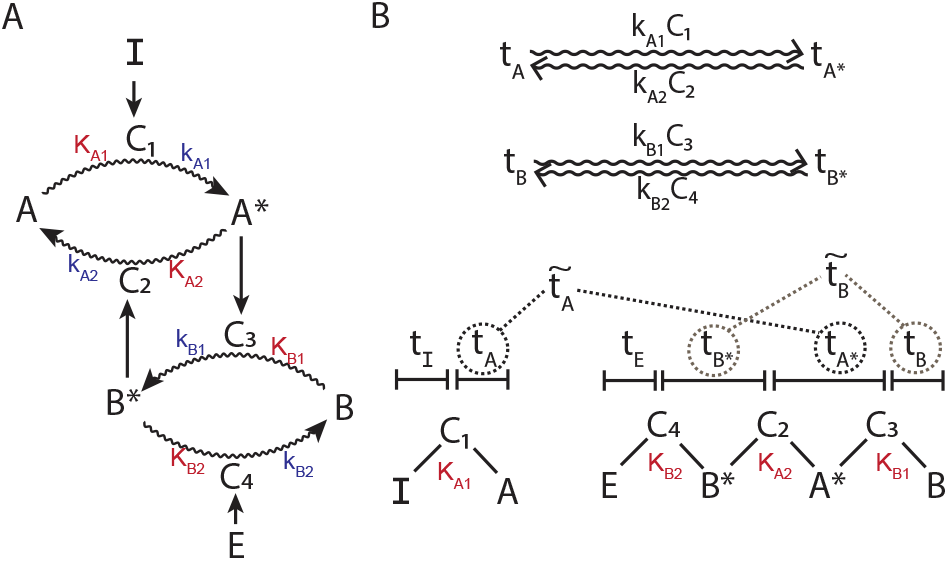
Enzymatic negative-feedback loop and the competitive binding network analyzed here.

**Fig. 9.**
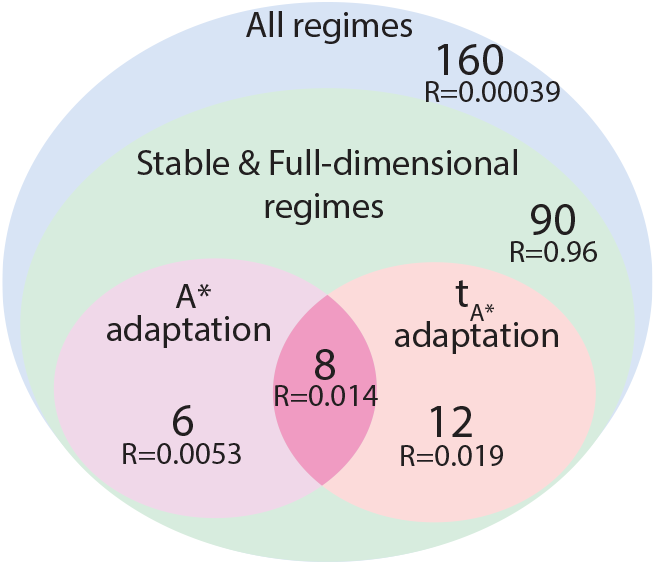
Classification of competitive-binding regimes that realize *A*^∗^ and/or 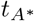 adaptation.

The slow dynamics are

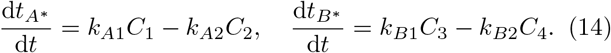

The classical reduced analysis uses Michaelis-Menten forms for the four complexes and obtains adaptation when 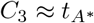 and *C*_4_ ≈ *t*_*E*_, so that

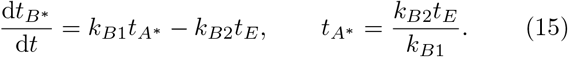

This expression is independent of *t*_*I*_ and therefore gives steady-state invariance.

Recent work questioned whether this conclusion survives competitive binding, where *A*^∗^ can bind either *B* or *B*^∗^ but not both (Jeynes-Smith and Araujo, 2023). We therefore analyzed the competitive binding network

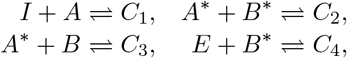

with binding constants *K*_*A*1_, *K*_*A*2_, *K*_*B*1_, *K*_*B*2_ and the corresponding total-concentration constraints

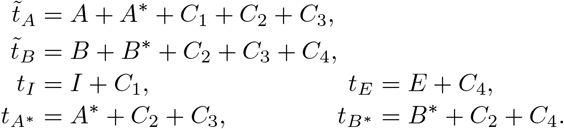

This network has a total of 276 dominance regimes. We look for stable fixed points of the catalysis dynamics. Therefore we impose (1) flux balance, and (2) local stability conditions on each regime. Also, we look for adaptation, requiring (3) steady-state inavriance with respect to *t*_*I*_, and (4) input responsiveness. Lastly, we are asking about *R*-index, therefore the regimes of interest should have a positive *R*-index, requiring its regime polyhedron to be (5) full-dimensional in parameter space.

The resulting valid regimes are summarized in Table 5. Competitive binding does not eliminate adaptation: 14 regimes realize free-*A*^∗^ adaptation, 20 realize 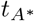 adaptation, and 8 regimes are shared. The resulting *R*-indices are *R*(*A*^∗^) = 0.0194±0.00004 and 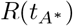 = 0.03291±0.00005.

**Table 5.** Regime counts and *R*-indices for competitive negative-feedback adaptation.

| Output | Valid regimes | Shared regimes | $R$-index |
| --- | --- | --- | --- |
| $A^*$ | 14 | 8 | $0.0194 \pm 0.00004$ |
| $t_{A^*}$ | 20 | 8 | $0.03291 \pm 0.00005$ |

The largest shared regime has dominance relations

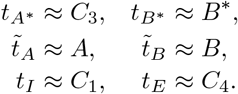

and reduced dynamics

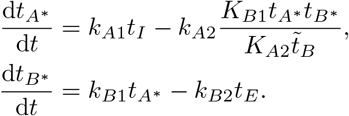

At steady state,

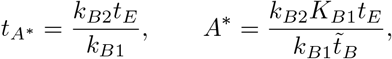

so both outputs are independent of *t*_*I*_.

Note that, similar to Eq. 15 from (Ma et al., 2009), this regime also has *t*_*E*_ dominated by *C*_4_, and 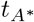 dominated by *C*_3_, so 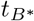 has the same reduced dynamics. However, the degradation rate of 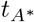 does not follow Michaelis-Menten form, therefore is excluded from Ma et al. (2009)’s analysis. However, this regime shares the same control mechanism of adaptation, namely that that 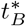 acts as an integral variable of the “error” of adaptation of 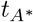, forming an integral negative feedback loop.

This provides an explicit regime-level explanation for adaptation under competitive binding. A numerical simulation from this regime shows transient responses after input changes followed by recovery of both outputs (Fig. 10).

**Fig. 10.**
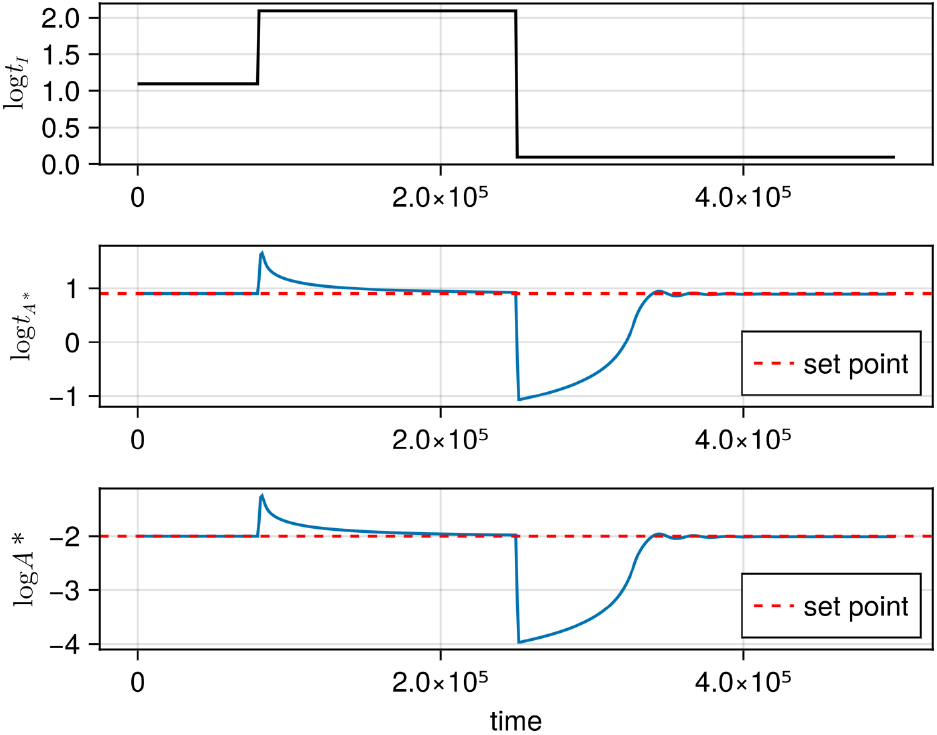
Simulated adaptation of *A*^∗^ and 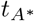 after perturbations of *t*_*I*_. With parameters: 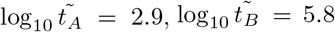, log_10_ *t*_*E*_ = −1.3, log_10_ *K*_*A*1_ = −1.1, log_10_ *K*_*A*2_ = 2.9, log_10_ *K*_*B*1_ = 2.9, log_10_ *K*_*B*2_ = 0, log_10_ *k*_*A*1_ = −3.4, log_10_ *k*_*A*2_ = 0, log_10_ *k*_*B*1_ = −2.2, log_10_ *k*_*B*2_ = 0, log_10_ *t*_*I*_ start with 1.1, change to 2.1 at *t* = 8, 000 and change to 0.1 at *t* = 25, 000.

This analysis highlights that numerical parameter sweeps, as used in (Jeynes-Smith and Araujo, 2023), could easily miss important behaviors when dimensionality slightly increases, and lead to incorrect conclusions. Under the parameter box used in (Jeynes-Smith and Araujo, 2023), namely 10^0^–10^4^ for protein concentrations and 10^−3^–10^4^ for binding and catalytic constants, the constrained *R*-index in this case for *A*^∗^ adaptation shrinks to about 3 × 10^−7^. This makes it hard to find using numerical scans of millions of points. This also highlights the importance of a definition of *R*-index that is unbiased by considerations of specific scenarios, so that behaviors in the network can be holistically explored and evaluated.

## 5. DISCUSSION AND DESIGN IMPLICATIONS

The examples above illustrate three practical uses of the *R*-index. First, it separates a reduced model from the conditions that make the model valid. The Michaelis-Menten case shows that the familiar textbook assumption *t*_*S*_ ≫ *t*_*E*_ is sufficient but not necessary; a distinct region, *K*_*m*_ ≫ *t*_*E*_, also supports the same reduced expression. Thus regime analysis can recover hidden validity regions missed by deriving the approximation from one intuitive limiting argument.

Second, the *R*-index makes explicit the dependence of realizability on the choice of control variables. A biocircuit may be operated at a single point, over an input range, or under a constrained experimental protocol. These scenarios correspond to different projections of the regime polyhedra and can produce different realizability scores. In this sense, the *R*-index, and the realizability and biological relevance implied by it, is not a property of a biochemical topology, but of a topology together with a design task.

Third, the method provides a route from qualitative design rules to quantitative comparisons. For example, approximate equality between independent parameters has zero asymptotic volume, but can become robust if enforced by circuit architecture, shared binding domains, or common degradation mechanisms. Conversely, a positive unconstrained *R*-index can become very small after experimental bounds are imposed, as in the competitive adaptation example. These observations suggest that high-*R* designs should be sought by choosing topologies with large valid regime cones or by engineering dependencies that reduce the effective dimension of parameter space.

The current calculation is asymptotic and log-geometric. It therefore complements, rather than replaces, numerical simulations and parameter fitting. Simulations test behaviors at sampled points; the *R*-index describes the size and structure of the regions from which those points are drawn. Combining the two can make numerical screens more informative: sampling can be biased toward candidate regime cones, and failed screens can be interpreted as evidence about the chosen parameter box rather than only the circuit topology.

Several implementation details also matter. The present calculations use asymptotic inequalities such as *A* ≫ *B*, so the numerical threshold chosen by an experimentalist, for example ten-fold or hundred-fold separation, will affect finite-box estimates. The solid-angle definition avoids dependence on any one finite threshold, but constrained estimates should use thresholds and parameter ranges matching the intended biological platform. Likewise, parameters that are biophysically linked should not be sampled independently. Shared domains, conserved catalytic motifs, or common expression systems can create correlations that change the effective geometry of the valid region.

Another advantage of the regime representation is interpretability. Each valid cone comes with dominance relations and reduced equations, so the *R*-index does more than rank circuit designs by realizability: it also explains which physical mechanism produces the desired behavior. In the adaptation example, valid competitive-binding regimes are clarified by the dominance *t*_*E*_ ≈ *C*_4_ and the integral-like equation for 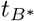. This mechanistic explanation is harder to extract from a parameter screen alone, especially when the successful parameter fraction is below one percent.

Finally, the same workflow can be used prospectively. Given a library of candidate network topologies, one can compute valid regime cones for each topology, compare their *R*-indices under the same control-variable assumptions, and prioritize the designs with the largest or most experimentally accessible cones. This suggests a design loop in which topology search, parameter-range specification, and experimental implementation are evaluated in a common geometric language.

## 6. SUMMARY

The *R*-index quantifies how much of log-parameter space supports a desired biocircuit function. Applied to standard formula of bioregulation such as Michalies-Menten and Hill, it distinguishes sufficient textbook conditions from necessary regime-level validity conditions and shows how realizability depends on which variables are experimentally controlled. For example, Hill function’s *R*-index goes to zero as the Hill coefficient grows when total substrate is free to vary. For competitive enzymatic negative feedback, *R*-index analysis reveals adaptation of both *A*^∗^ and 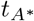 are possible, and finds explicit regimes revealing mechanisms of adaptation. While resolving an active debate in the field, this highlights the shortcomings of numerical parameter sweeps in high-dimensional spaces, as well as the importance of holistic analysis provided by the *R*-index.

These results suggest that regime-based analysis that *R*-index uses can serve as a validity-aware design language: instead of asking only whether a reduced model can be written, it asks where the model is valid and how difficult that region is to realize experimentally. This has promising applications in studying and designing functional biocircuits.

